# Dual-species proteomics and targeted intervention of animal-pathogen interactions

**DOI:** 10.1101/2023.07.02.547426

**Authors:** Yang Sylvia Liu, Chengqian Zhang, Bee Luan Khoo, Piliang Hao, Song Lin Chua

**Affiliations:** Department of Applied Biology and Chemical Technology, The Hong Kong Polytechnic University, Kowloon, Hong Kong SAR China; School of Life Science and Technology, ShanghaiTech University, China; Department of Biomedical Engineering, City University of Hong Kong, Hong Kong SAR China; Hong Kong Center for Cerebro-Cardiovascular Health Engineering (COCHE); City University of Hong Kong-Shenzhen Futian Research Institute, Shenzhen; State Key Laboratory of Chemical Biology and Drug Discovery, The Hong Kong Polytechnic University, Kowloon, Hong Kong SAR China; Research Centre for Deep Space Explorations (RCDSE), The Hong Kong Polytechnic University, Kowloon, Hong Kong SAR China

**Keywords:** SILAC Proteomics, *Pseudomonas aeruginosa*, *Caenorhabditis elegans*, host-pathogen interactions

## Abstract

Complexity in host-pathogen interactions drives the need to develop sensitive and accurate biochemical techniques to elucidate host and pathogen protein expressions. Current proteomics techniques reveal information from the point of view of either the host or pathogen, but do not provide data on the corresponding partner. While dual-species transcriptomics is increasingly used to study RNA expression in host and pathogen, it remains challenging to simultaneously study host-pathogen proteomes that reflect the direct competition between host and pathogen. Using *Caenorhabditis elegans*-*Pseudomonas aeruginosa* infection model as proof-of-concept, we established a forward+reverse SILAC proteomics approach to simultaneously label and quantify newly-expressed proteins of host and pathogen without physical isolation. We observed iron competition between pathogen iron scavenger and host iron uptake protein, where *P. aeruginosa* upregulated pyoverdine synthesis protein (PvdA) and secreted pyoverdine, and *C. elegans* expressed ferritin (FTN-2) respectively. Using Galangin as a novel PvdA inhibitor identified by structure-based virtual-screening, targeted intervention of iron competition eliminated *P. aeruginosa* infection, and enabled animal survival. Our work provides insights into the mechanisms dictating host-pathogen interactions and offers novel strategies for anti-infective therapy.

## Introduction

Systems biology approaches are increasingly employed to study host-pathogen interactions, because of the technical advances in specificity and sensitivity. Quantitative proteomics are used widely to study either pathogen or host proteomics in an infection (1, 2), but it is important to note that proteomics was typically performed only on the host after treatment with microbial proteins or microbe itself (from the point of view of the host) (3, 4), or on the microbe after interactions with the host (from the point of view of microbe) (5, 6). This implied that researchers can only acquire one-sided proteomic data from either the host or microbe, without gaining proteomic perspectives from the other partner. In an effort to elucidate novel host-pathogen interactions, host and pathogen co-study using proteomics is gaining traction in recent years (7, 8). Ideally, dual-species proteomics quantifies proteins of pathogens and animal host in a single experiment, and provides insights into both host and pathogen responses during infection. However, many strategies require pre-isolation steps which are destructive and difficult to study small bacterial populations in the larger host (5).

One accurate and sensitive proteomics technique frequently used to accurately label eukaryotic proteomes for liquid chromatography-mass spectrometry (LC-MS) analysis is stable Isotope Labeling with Amino acids in Cell culture (SILAC) (9). However, its application in studying both animal and pathogen during infections is limited, due to difficulty in labeling both entire animal and prokaryotic pathogens. This provides a rationale for us to develop a SILAC-based proteomics approach to simultaneously characterize host and pathogen physiologies during infection.

Bacterial pathogens establish chronic infections by forming biofilms on host tissues and medical implants (10), where 80% bacterial infections attributed to biofilms (11). Biofilms are bacterial aggregates embedded in protective exopolymeric substances (EPS) (12), allowing bacteria to attain extreme resistance to antimicrobials and host clearance (13, 14). As model organism for biofilm research and member of the dangerous ESKAPE pathogens (15), *Pseudomonas aeruginosa* is a Gram negative pathogen that causes infections in hospitalized patients, immunocompromised individuals and elderly, where mortality rate for bacteremia can reach 39% (16, 17). *P. aeruginosa* could infect the respiratory system (18), and the intestinal tract, such as gut-derived sepsis, Shanghai fever and severe necrotizing enterocolitis (19–22). The intestinal tract is also an important reservoir for *P. aeruginosa* opportunistic infections and dissemination to the lungs (23). Moreover, *P. aeruginosa* infections could range from localized infections of the skin to life-threatening systemic diseases or seed following bloodstream infections (24, 25).

*P. aeruginosa* employs a plethora of virulence factors for establishing biofilm-mediated infections. Its ability to produce the sticky biofilm matrix, which comprises of exopolysaccharides, adhesion proteins and extracellular DNA (eDNA), to prevent antibiotics and immune cells from accessing the biofilm cells (26). Another virulence factor is pyoverdine, which is a secreted self-fluorescent siderophore important for uptake of iron from the environment and biofilm formation (27). Biofilm formation and pyoverdine production are mainly attributed to c-di-GMP secondary messenger signaling, which is ubiquitous in many bacterial species (28, 29).

To study bacterial virulence of intestine infection, *Caenorhabditis elegans* is commonly used as an experimental host to study host-pathogen interactions, because of its similar mechanisms of immune defense with orthologs in mammals and invertebrates (30). Due to these universal host responses, microbial pathogenesis has evolved correspondingly, where bacterial pathogens share similar host-infection strategies. For instance, *C. elegans* upregulated immune response genes, such as autophagy pathway and lysozymes, while different pathogens upregulated host invasion proteins (31–34). Development of anti-virulence agents that interfere host-pathogen interactions will tilt the balance in favor of the host (35), leading to the elimination of pathogens and recovery from infectious diseases.

Iron is a vital nutrient for both host and pathogen, but its availability is often limited in the host’s extracellular environment due to sequestration mechanisms. As a result, host-pathogen interactions are often characterized by intense competition for this essential nutrient (36–38). In response to this competition, both hosts and pathogens have evolved various strategies to acquire iron. For instance, pathogens secrete iron-chelating siderophores such as staphyloferrin in *Staphyloccoccus aureus* (39), while hosts produce iron uptake proteins such as lactoferrin. The identification of strategies used by both host and pathogen to compete for iron has led to the development of novel anti-virulence therapies that target these pathways, potentially reducing pathogen virulence and improving host outcomes.

Using a *C. elegans* host-*P. aeruginosa* pathogen interaction model as proof-of-concept, we established a concurrent forward (host)-reverse (microbe) SILAC proteomics approach to label and quantify newly synthesized host and pathogen proteins after *P. aeruginosa* establish infections in the *C. elegans* intestine. For the host, its ^12^C light lysine-proteins will incorporate ^13^C heavy lysine in its newly synthesized proteins, while the^13^C heavy lysine-labeled pathogen will correspondingly incorporate ^12^C-_L_-lysine into its proteins during host infection. This represents a technical advance over existing proteomics techniques that only evaluate either host or microbe proteomes.

Specifically, our proteomics data revealed that *P. aeruginosa* upregulated its expression of pyoverdine synthesis proteins (PvdA), whereas *C. elegans* expressed the ferritin protein (FTN-2), indicating that both host and pathogen were using iron scavenging strategies in iron competition. Although a previous study had shown that FTN-2 was part of the immune network upregulated in response to *S. aureus* (*40*), we showed for the first time that pyoverdine was the direct iron competitor of FTN-2.

To interfere effective iron competition by the pathogen, we employed structure-based virtual screening to identify Galangin as a novel *pvdA* inhibitor that can reduce pyoverdine expression. The Galangin-ciprofloxacin combinatorial therapy could eliminate *P. aeruginosa* infection and promote host survival. Hence, our proof-of-concept study developed the first SILAC proteomics approach which can simultaneously provide proteomic information of both host and pathogen in the same sample, thereby providing direct information about true host-pathogen interactions and novel insights into the development of anti-virulence strategies that intervene host-pathogen interactions.

## Results

### Using forward-reverse SILAC to study animal-pathogen interactions

As proof-of-concept, we employed a *C. elegans*-*P. aeruginosa* infection model by feeding the nematodes with *P. aeruginosa* biofilms. We had previously shown that *P. aeruginosa* formed biofilms on the agar (41, 42). We monitored the feeding of *P. aeruginosa* into *C. elegans* intestines after 6 hrs post infection (h.p.i) (early feeding) and 24 h.p.i. (late feeding), where there was increase in viable bacterial counts (**Fig. 1A**) and live *gfp*-tagged bacteria within the intestine (**Fig. 1B**). This indicated that *P. aeruginosa* colonized and remained viable in the *C. elegans* intestine, while its GFP production indicated that they were metabolically active.

**Figure 1:**
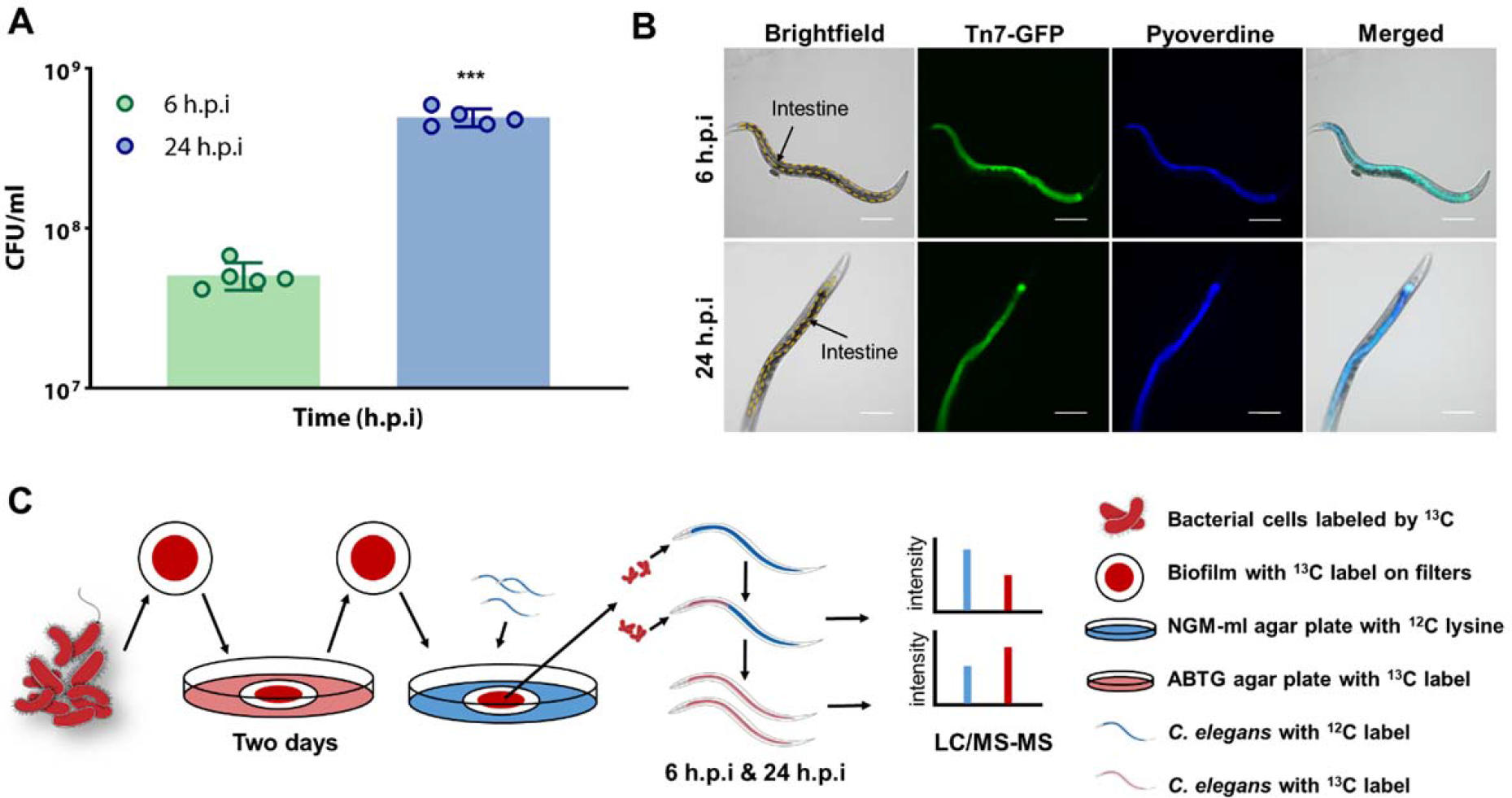
Workflow of SILAC proteome analysis of *C. elegans-P. aeruginosa* host-pathogen interaction model. (**A**) *P. aeruginosa* population numbers in intestine at 6 h.p.i and 24 h.p.i. (**B**) Fixed *C. elegans* with *gfp*-tagged PAO1 in intestine at 6 h.p.i and 24 h.p.i. (**C**) The workflow and experimental design of this project, including the *P. aeruginosa – C. elegans* infection model and the SILAC proteomics assay. Δ*lysA* mutant labeled with ^13^C-_L_-lysine was cultivated on agars which contained ^13^C-_L_-lysine for two days to form biofilm. The samples were then collected for the SILAC proteomics assay. H.p.i = hours post-infection. Scale bar: 100 μm. ***P < 0.001.

To understand the overall interactions between *C. elegans* and intestinal *P. aeruginosa*, we modified and established a forward-reverse SILAC workflow (**Fig. 1C**), which could study new protein abundances in the intestinal *P aeruginosa* and animal host quickly and accurately by labeling host and bacterial proteins simultaneously without the need for physical isolation. The SILAC proteomic approach is commonly used to determine changes in protein expression induced by experimental conditions in mammalian cell culture (43, 44), with few applications in prokaryotes (45–47). Our SILAC approach broadened the scope of SILAC proteomics, where mechanisms of host-pathogen interactions could be identified and applied to other bacterial species.

To ensure that *P. aeruginosa* could only incorporate _L_-lysine for SILAC labelling, we employed the *P*. *aeruginosa* mutant that cannot self-synthesize lysine, mPAO1Δ*lysA* and requires exogenous lysine to survive (48). We previously showed that the Δ*lysA* mutant with _L_-lysine supplements had similar growth rates and metabolism as wild-type PAO1 (48). Within the context of this study, we also showed that as long as the medium was supplemented with _L_-lysine, the Δ*lysA* mutant could colonize and infect *C. elegans* intestine, and kill the animal over time in a similar fashion as wild-type PAO1 (**Supplementary Figure 1**). As for *C. elegans*, lysine is a dietary essential amino acid (49), with 93% incorporation of labelled lysine into its proteins (50), so the animal could be directly used for our proteomic approach.

Hence, the Δ*lysA* mutant prelabeled with ‘heavier’ ^13^C-_L_-lysine were initially fed to *C. elegans* whose proteins contain ‘lighter’ ^12^C-_L_-lysine. Over the course of host-microbial interactions, intestinal *P. aeruginosa* could incorporate ^12^C-_L_-lysine from the host in its newly synthesized proteins, while *C. elegans* would instead incorporate ^13^C-_L_-lysine e from intestinal *P. aeruginosa* in its new proteins. The Q-Exactive HF-X Hybrid Quadrupole-Orbirap Mass spectrometer system was employed to analyze the SILAC samples, where we identified more than 4000 proteins differentially expressed by *C. elegans* and *P. aeruginosa* from established databases (WormBase for *C. elegans* (51) and Pseudomonas.com for *P. aeruginosa* (52)) using the Proteome Discoverer software (PD) (53). Only proteins that were differentially expressed at higher (>2-fold) or lower (<0.5-fold) levels in triplicate samples at 24 h.p.i than 6 h.p.i were shortlisted for further analysis.

We hypothesized that the actively expressed proteins by the host and pathogen are important in conferring competitive advantages over the other. A total of 752 proteins from *P. aeruginosa* and 3386 proteins from *C. elegans* were subjected to MetaboAnalyst 5.0 for statistical analysis (54). For the multivariate analysis of *P. aeruginosa* and *C. elegans* proteins, the principal component analysis (PCA) plot (**Fig. 2A-B**) revealed that the 6 h.p.i group and 24 h.p.i group were clustered separately. The PCA plot of *P. aeruginosa* was performed with the 61.9% of total variance between these two groups represented by the first two principal components (**Fig. 2A**). The principal component 1 (PC1) and principal component 2 (PC2) explained 43% and 18.9% of the variance (**Fig. 2A**). For multivariate analysis of *C. elegans* proteins, the principal component analysis (PCA) plot revealed that two groups were with the 85% of total variance represented by the first two principal components (**Fig. 2B**). The principal component 1 (PC1) and principal component 2 (PC2) explained 72.7% and 12.3% of the variance (**Fig. 2B**). The volcano plot combines a measure of statistical significance from statistical analysis with the magnitude changes (55). These dots (blue: downregulated, red: upregulated) in volcano plot of *P. aeruginosa* (**Fig. 2C**) and *C. elegans* (**Fig. 2D**) indicates different proteins that display both large magnitude fold-changes (x axis) and high statistical significance (-log10 of p values, y axis).

**Figure 2:**
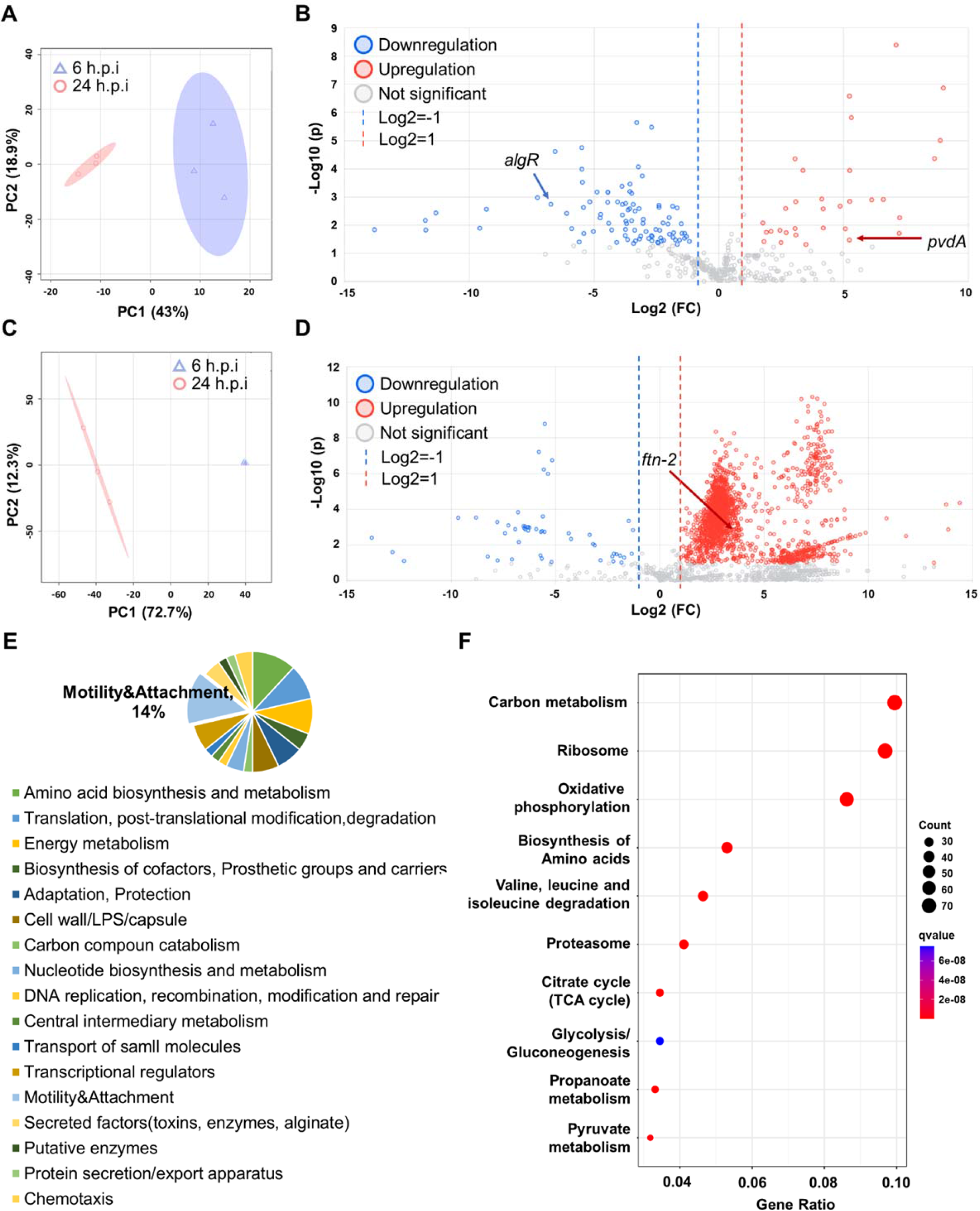
Proteomic profiling of proteins in *P. aeruginosa* and *C. elegans* during interactions. (**A**) The PCA score plots showed that the proteins of *P. aeruginosa* samples from 6 h.p.i and 24 h.p.i groups were clustered separately. (**B**) Volcano plot representation of differential expression analysis of proteins in *P. aeruginosa* samples from 6 h.p.i and 24 h.p.i groups. (**C**) The PCA score plots showed that the proteins of *C. elegans* samples from 6 h.p.i and 24 h.p.i groups were clustered separately. (**D**) Volcano plot representation of differential expression analysis of proteins in *C. elegans* from 6 h.p.i and 24 h.p.i groups. (**E**) Functional groups of upregulated proteins of *P. aeruginosa* from 6 h.p.i to 24 h.p.i. The Motility & Attachment took a large proportion in upregulated groups. (**F**) Significantly enriched KEGG pathways of the upregulated proteins from *C. elegans*.

Protein AlgR (gene=*algR*) and PvdA (gene=*pvdA*) of *P. aeruginosa* were downregulated and upregulated with a log2 (FC) value of -6.7367 (pvalue=0.0018066) and 5.2313 (pvalue= 0.033246) respectively (**Fig. 2C**). Protein FTN-2 of C. elegans was upregulated with a log2 (FC) value of 3.4057 (pvalue= 0.0016067) (**Fig. 2D**). According to the standards of Functional Classifications Manually Assigned by PseudoCAP for bacteria(56), the upregulated and downregulated proteins of *P. aeruginosa* were classified into different functional groups (**Fig. 2E** and **Supplementary figure 2**).

Our initial analysis revealed that for *P. aeruginosa*, a large proportion (14%) of motility and attachment proteins were upregulated from 6 h.p.i to 24 h.p.i (**Fig. 2E**), which corroborated with previous findings that motility and surface attachment are important in bacterial colonization in the host (57). After uploading proteins information of *C. elegans* to KEGG database, the pathway enrichment of proteins set was analyzed through clusterProfiler package in R. From the gene ratio in the dot plot (**Fig. 2F**), the most enriched KEGG pathways were carbon metabolism, ribosome and oxidative phosphorylation, which were similar to previous study (58). According to the WormBase gene ontology for *C. elegans*, upregulated and downregulated proteins from host were functionally grouped in **Supplementary figure 3** respectively. This indicated that our SILAC approach was reliable in detecting changes in protein expression accurately.

Interestingly, among bacterial proteins, we observed that the positive alginate biosynthesis regulatory protein AlgR(59) was significantly downregulated, while L-ornithine N(5)-monooxygenase PvdA(60) was upregulated from 6 h.p.i to 24 h.p.i. AlgR is a regulator involved in the expression of multiple virulence factors, such as pyoverdine and other iron acquisition genes (PvdS) (61, 62), whereas PvdA is involved in the biosynthesis of pyoverdine, an iron siderophore important for microbial growth (63). Correspondingly, *C. elegans* upregulated the expression of FTN-2 (ferritin), an iron scavenging protein commonly found in most eukaryotes (64). Since both host and pathogen upregulated their mechanisms important in iron uptake, this raised the rationale that the pathogen and animal used pyoverdine and ferritin respectively to compete for iron. While pyoverdine was recently implicated in *C. elegans* intestinal infection (65) and ferritin was involved in innate immune response (66), it was unclear if both were direct iron competitors in *C. elegans* infection.

### Downregulation of algR promotes pyoverdine production in C. elegans

Since our proteomics data revealed a correlative downregulation of AlgR and PvdA upregulation, we next aim to show that intestinal *P. aeruginosa* had lower AlgR expression to increase PvdA expression within *C. elegans* intestine, where pyoverdine was the virulence factor that caused *C. elegans* death. The *pvdA* gene expression (67) and corresponding pyoverdine production was upregulated in intestinal wild-type PAO1 within *C. elegans* from 6 h.p.i to 24 h.p.i, whereas the Δ*pvdA* mutant did not (**Fig. 3A-B**). Hence, we observed that the Δ*pvdA* mutant lost its ability to kill *C. elegans*, which can be restored with the *pvdA* complementation (**Fig. 3C**). The expression of *pvdA* was inversely regulated by *algR* gene. The Δ*algR* mutant displayed higher expression of *pvdA* gene and pyoverdine levels than wild-type PAO1 and Δ*pvdA* (**Fig. 3A-B**), which confirmed the findings from proteomic data. This had severe implications in the animal host, where the Δ*algR* mutant could kill *C. elegans* at a higher and faster rate than wild-type PAO1 (**Fig. 3C**). Gene complementation in Δ*algR* mutant restored inability by *P. aeruginosa* to effectively kill the populations (**Fig. 3C**). This corroborated with previous study that AlgR expression repressed pyoverdine production (68). Hence, this showed that the downregulation of AlgR resulted in the higher expression of pyoverdine, which was crucial in killing the host animals.

**Figure 3:**
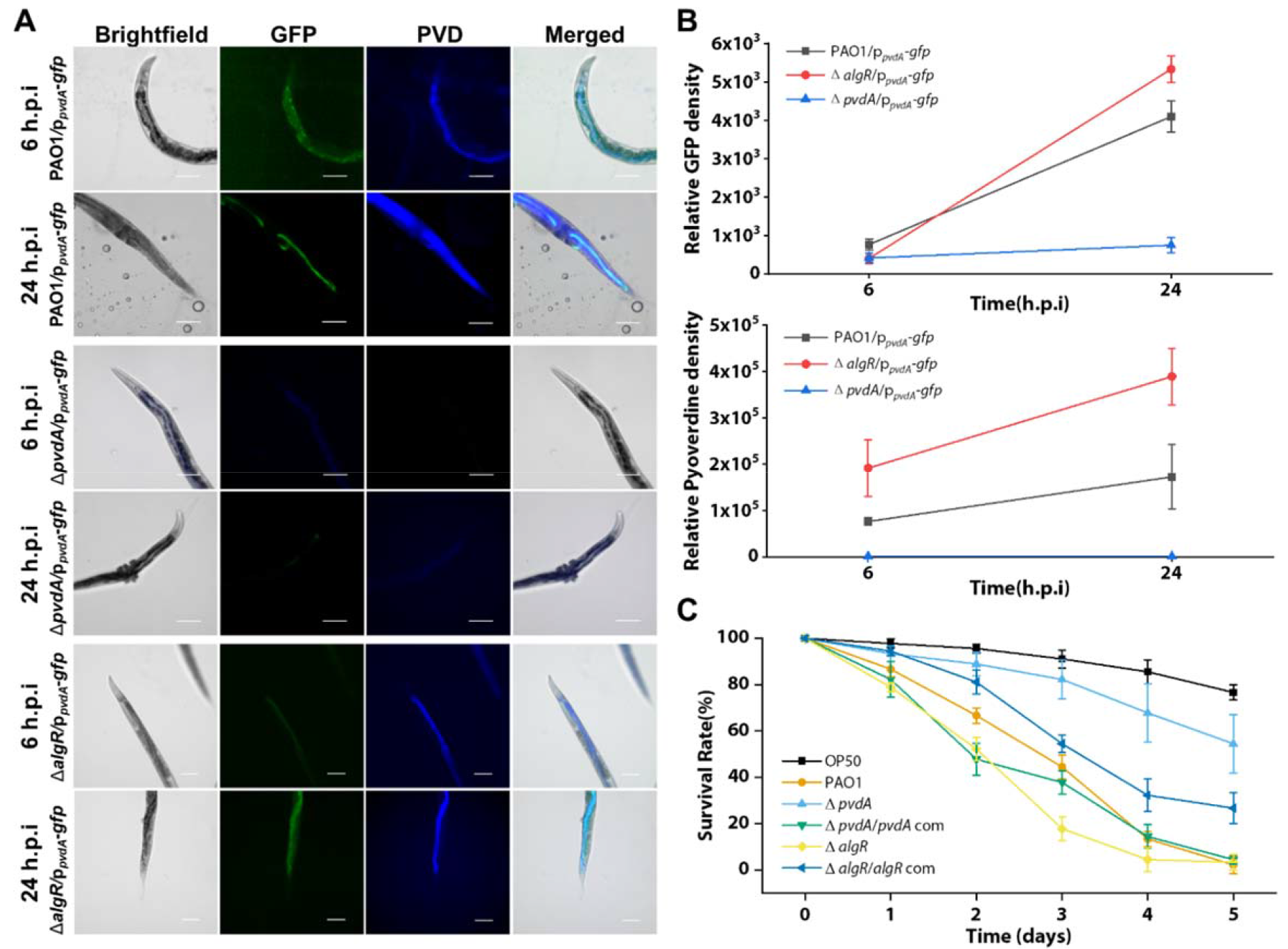
*P. aeruginosa* upregulates pyoverdine expression via repression of AlgR and enhancing of PvdA in intestinal infection of *C. elegans*. (**A**) Representative fluorescent images of the fixed nematode’s intestine after the infection of PAO1/p*_pvdA_-gfp*, Δ*pvdA*/p*_pvdA_-gfp* and Δ*algR*/p*_pvdA_-gfp* from 6^th^ to 24^th^ hour. GFP is green, and Pyoverdine channel is blue. Scale bars, 50 μm. (**B**) Relative GFP fluorescence density and pyoverdine density from wild-type (N2) nematodes infected by PAO1/p*_pvdA-_gfp*, Δ*pvdA*/p*_pvdA_-gfp* and Δ*algR*/p*_pvdA_-gfp* from 6^th^ to 24^th^ hour. (**C**) Nematodes were cultivated on different bacterial mutants’ lawn for 5 days. The survival rate of nematodes was recorded. H.p.i = hours post-infection. Scale bar: 100 μm.

### Host ferritin is involved in iron competition with bacterial pyoverdine

From the point of view of *C. elegans* host, our proteomics data revealed the upregulation of FTN-2 protein which was involved in iron metabolism (69). By using the *ftn-2*(ok404) mutant animal (RB668) deficient in ferritin production, we surprisingly observed that it did not elicit the iron scavenging response from *P. aeruginosa*. In wild-type animals, *P. aeruginosa* upregulated *pvdA* gene expression and pyoverdine production (**Fig. 4A-C**). However, there was subdued *pvdA* expression and lower associated pyoverdine levels by intestinal *P. aeruginosa* when colonizing *ftn-2*(ok404) mutant animal (**Fig. 4A-C**). Interestingly, the *ftn-2*(ok404) mutant animals could survive better than wild-type N2 animals upon *P. aeruginosa* infection (**Fig. 4C**). This indicated that without the host iron competitor (FTN-2), the intestinal bacteria did not seem to activate its pyoverdine operon, thereby negating the iron competition between host and pathogen.

**Figure 4:**
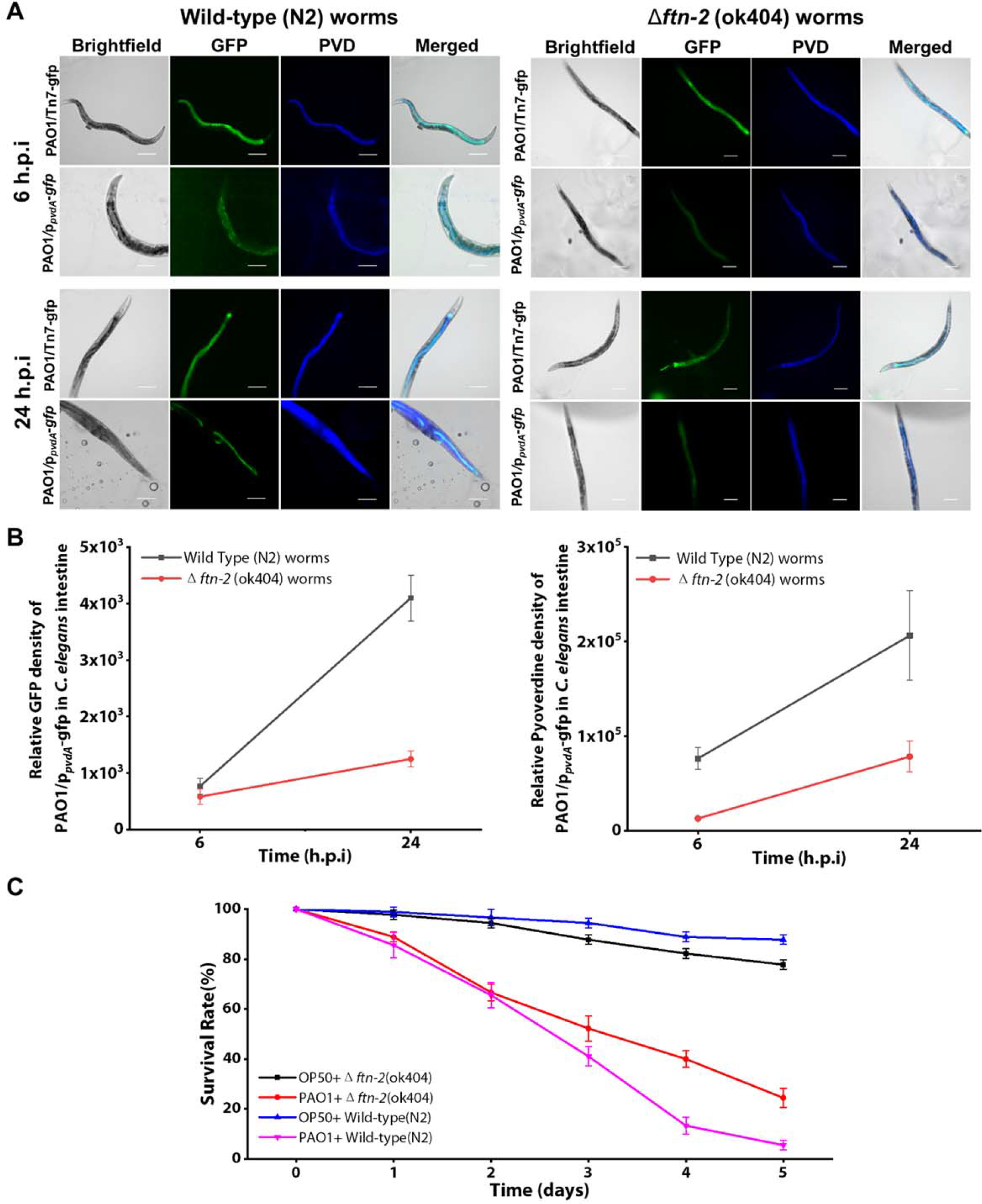
Role of *FTN-2* by *C. elegans* in iron competition during the bacterial infection interaction compared to wild type nematodes. (**A**) Representative fluorescent images of the fixed wild type (N2) and *ftn-2* (ok404) nematodes intestine after the infection of PAO1/Tn7-gfp and PAO1/p*_pvdA_-gfp* from two time points (6 h.p.i and 24 h.p.i). (**B**) Relative GFP density and Pyoverdine density of the nematodes intestinal PAO1/p*_pvdA_-gfp* biosensor between wild type nematodes and *ftn-2* (ok404) mutant nematodes over time. Both GFP promoted by PvdA and pyoverdine of *P. aeruginosa* had high expression in wild-type *C. elegans* intestine than *ftn-2* (ok404) *C. elegans*. (**C**) The survivability of wild-type *C. elegans* compared to *ftn-2* (ok404) *C. elegans* infected by PAO1 for 5 days. Compared to the wild type nematodes, *ftn-2* (ok404) nematodes had less competition with bacteria. H.p.i = hours post-infection. Scale bar: 100 μm.

### Galangin is a pvdA inhibitor that interferes host-pathogen interactions

To intervene iron competition between host and pathogen and tile the favor towards the host, we next aim to develop an anti-virulence strategy that inhibits PvdA and intercept pyoverdine synthesis in *P. aeruginosa*. While iron chelators such as EDTA and DIPY were previously proposed as anti-pyoverdine strategies (70, 71), these chemicals may have pleiotropic effects, thereby requiring inhibitors specific against pyoverdine synthesis proteins. Another inhibitor (5-fluorocytosine) also targeted PvdS, regulator in the *pvd* synthesis pathway (72).

We employed structure-based virtual screening to identify potential inhibitors from the natural product compounds library that competitively bind to the active site of PvdA protein (73). We preliminary identified Galangin (3,5,7-trihydroxyflavone), a flavonoid which could bind to the PvdA active sites in a similar position as the N∼5∼-hydroxy-L-ornithine (ONH) ligand at stronger binding affinity (-6.7 kcal/mol for Galangin as compared to -4.8 kcal/mol for ligand) (**Fig. 5A**). We confirmed Galangin’s inhibition of PvdA by showing the inhibition of *pvdA* gene expression (**Fig. 5B**) and pyoverdine levels (**Fig. 5C**), where the half-maximal inhibitory concentration (IC50) is 22.69 μM (**Fig. 5D**).

**Figure 5:**
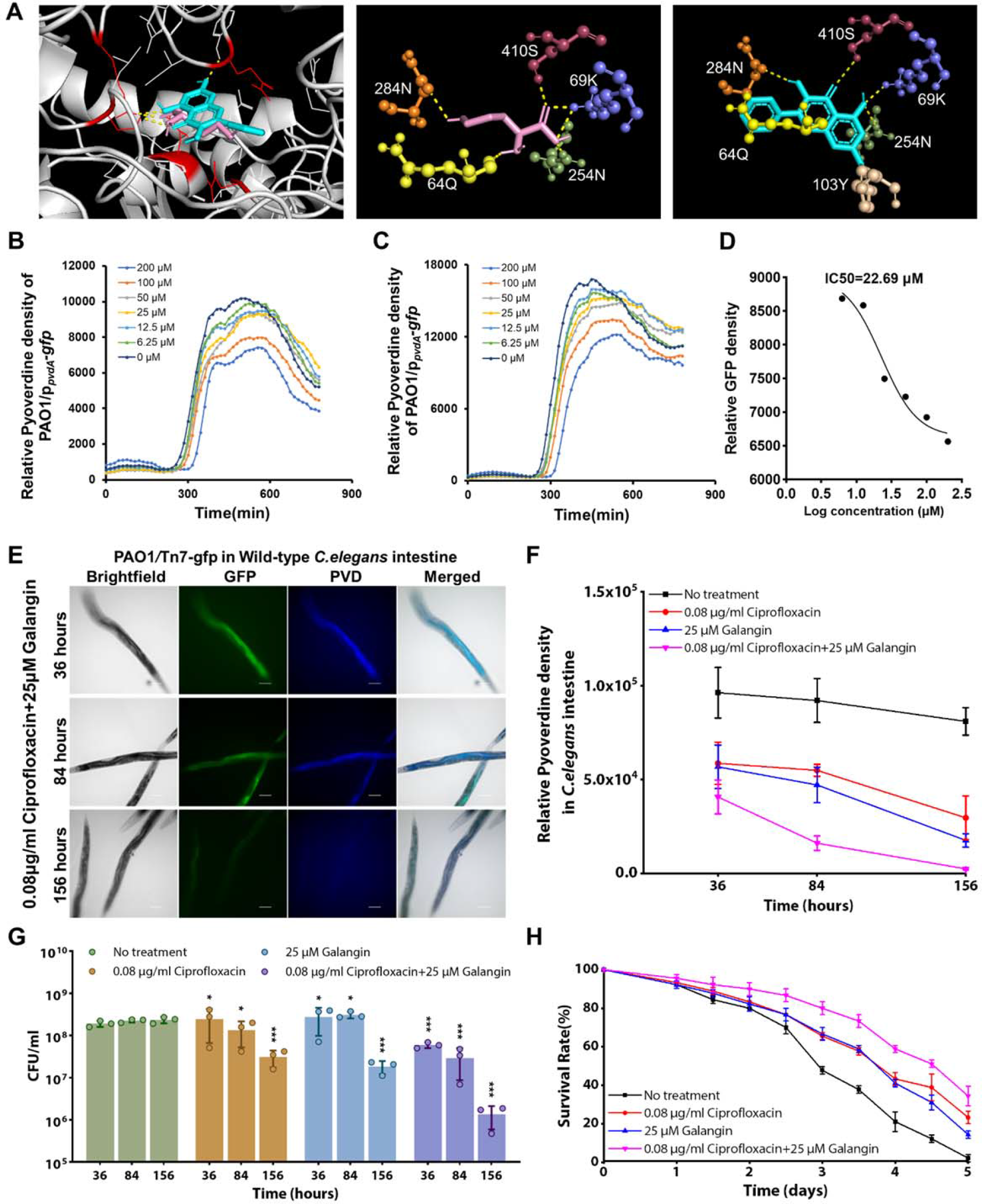
Galangin serves as PvdA inhibitor that abrogates P. aeruginosa intestinal infection. (**A**) Molecular docking revealed Galangin (blue) and ONH (pink) binding to amino acid residues in active sites of PvdA. (**B**) Galangin inhibits *in vitro pvdA* gene expression by testing GFP level of PAO1/p*_pvdA_-gfp*. (**C**) Galangin inhibits *in vitro* pyoverdine production by testing fluorescent density. (**D**) The half-maximal inhibitory concentration (IC50) graph of Galangin to *pvdA* expression and pyoverdine production inhibition. (**E**) Representative fluorescent images of fixed *C. elegans* intestine with Galangin and Ciprofloxacin treatment. (**F**) Relative pyoverdine density from wild type nematodes infected by PAO1 and then with the combinatorial treatment of Galangin and Ciprofloxacin. This combinatorial treatment can help to eliminate the intestinal *P. aeruginosa* infection in *C. elegans* in 156 hours with several times of drug administration. (**G**) *P. aeruginosa* CFU in nematodes intestine after Galangin and/or Ciprofloxacin treatment for 5 days. Bacteria numbers show an alleviation of biofilm infection after combinatorial treatment. (**H**) Survival rate of wild type *C. elegans* with Galangin and Ciprofloxacin treatment. This combinatorial treatment could increase survival rate of host. Scale bar: 100 μm. ***P < 0.001, **P < 0.01, *P < 0.05.

Although PvdA inhibition of Galangin is not bactericidal (**Supplementary Fig. 4**), this might prevent the pathogen from establishing effective infection in the host, so its combinatorial therapy with an antibiotic was expected to enhance the killing of pathogen in disease treatment. As proof-of-concept, we employed ciprofloxacin, which is widely used for the treatment of biofilm infections caused by *P. aeruginosa* (74). In *in vitro* assays, ciprofloxacin could inhibit *P. aeruginosa* biofilm growth with an IC50 of 0.04 μg/ml, and the minimal concentration of Galangin in combination with ciprofloxacin that could inhibit bacterial biofilm growth was 12.5 μM and 0.04 μg/ml respectively from the CFU results (**Supplementary figure 5**). As the IC50 of ciprofloxacin in monotreatment to inhibit biofilm formation was 0.05279 μg/ml (**Supplementary figure 5**), the fractional inhibitory concentration (FIC) index value of combinatorial therapy of Galangin and ciprofloxacin to biofilm formation *in vitro* is around 1.0 (>0.5, <4.0), which indicates that Galangin and ciprofloxacin could be used in combinatorial therapy (75). Since higher doses of antimicrobials are typically expected for *in vivo* animal studies (76), we evaluated the efficacy of 25 μM Galangin-0.08 μg/ml ciprofloxacin combinatorial treatment against the *C. elegans* intestinal infection. Although Galangin and ciprofloxacin monotherapies could reduce pyoverdine levels in *C. elegans* (**Fig. 5E-F**), they had mild killing effect on intestinal *P. aeruginosa* populations (**Fig. 5G**). Instead, the Galangin-ciprofloxacin combinatorial treatment could significantly downregulate pyoverdine levels (**Fig. 5E-F**) and kill *P. aeruginosa* in the animal intestine (**Fig. 5G**), resulting in enhanced survival of the animals over time (**Fig. 5H**).

### Galangin-ciprofloxacin combinatorial therapy is effective against biofilm infection in a fish tail wound infection model

To further validate the combinatorial treatment of Galangin with ciprofloxacin to biofilm infection in vertebrate animal model, we then applied 25 μM Galangin-0.08 μg/ml ciprofloxacin combinatorial treatment against the Medaka fish wounds infection as described in the workflow (**Fig. 6A**). Medaka fish has transparent larvae fish bodies for description of infection establishment process via *in vivo* imaging techniques, and many research tools for studying gene functions can be utilized on Medaka (77, 78). The established Medaka-*P. aeruginosa* infection model on fish skin-wound provided a convenient, visible and cheap way to further validate antibacterial drugs (79).

**Figure 6:**
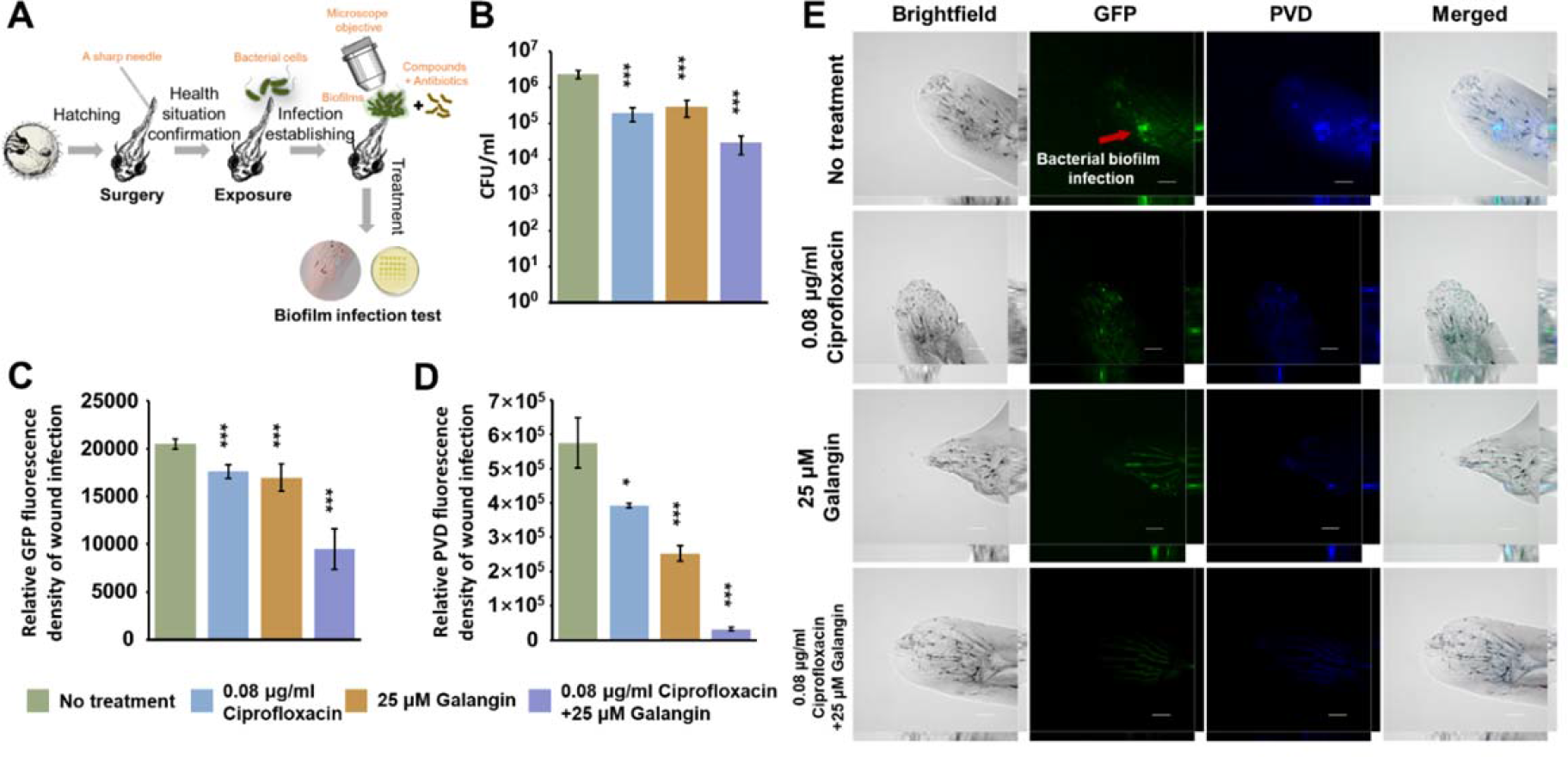
Combinatorial treatment of ciprofloxacin and Galangin eliminates the wounds infection on medaka fish tail wounds. (**A**) The establishment of medaka fish – *P. aeruginosa* infection model for testing novel antimicrobial strategies *in vivo*. (**B**) The bacterial populations in the wound infection after treatment by monotherapy and combinatorial therapy of Galangin and ciprofloxacin. (**C**) Relative GFP fluorescence expression level of PAO1/p*_pvdA_-gfp* on the wound infection after treatment by monotherapy and combinatorial therapy of Galangin and ciprofloxacin. (**D**) Pyoverdine expression level of in wound biofilm infection after monotherapy and combinatorial therapy of Galangin and ciprofloxacin. (**E**) Representative fluorescent images of wound bacterial (PAO1/p*_pvdA-_gfp*) biofilms infection on medaka fish after treatment. GFP: green, and pyoverdine: blue. Scale bar: 100 μm. The mean and ±s.d. from three experiments are shown. ***P < 0.001, **P < 0.01, *P < 0.05.

We observed that bacterial populations in the biofilm was reduced (**Fig. 6B**). The Galangin-ciprofloxacin combinatorial treatment could effectively downregulate the GFP expression level of p*_pvdA_-gfp* (**Fig. 6C**) and significantly lowered pyoverdine levels of wounds infection (**Fig. 6D**) from fluorescent images of treated fishes wound (**Fig. 6E**). Hence, the Galangin-ciprofloxacin combinatorial treatment is safe and effective in eliminating the wound infection in the fish. Hence, we demonstrated that Galangin could be used as PvdA inhibitor that intercept host-pathogen iron competition and tilt the survival advantage to the host.

## Discussion

Iron competition between host and pathogen has been studied extensively (36–38), but our study revealed the simultaneous upregulation of iron scavenging proteins from host and pathogen in the same animal infection model. This relied on the accuracy and sensitivity of our forward-reverse SILAC approach, which could achieve a reliable detection on proteins expression from both bacteria and host. Our method also represents a technical advance in proteomics which solve the existing problem of studying either the host or microbe proteome in the same sample. Although there are computational improvements where dual species can be possibly searched and identified in a single mass spectrometry run, the strength of our SILAC approach is its specificity and sensitivity, where the host has to incorporate the C-13-lysine from the bacteria directly and the bacteria can only incorporate C-12-lysine from the host. This ensures that any newly synthesized proteins must arise from direct interaction between host and pathogen, which is something that computational improvements cannot achieve. Furthermore, computational bias still exists in current models, where further normalization of samples is required.

Our proteomics data was further confirmed by our subsequent experiments showing that host or pathogen which was defective in iron scavenging could not participate in iron competition. Unexpectedly, *C. elegans* with ferritin mutation did not induce *P. aeruginosa* expression of pyoverdine, indicating that it ‘takes two hands to clap’ in host-pathogen interactions. Nonetheless, other virulence mechanisms could also come into play during the infection process, where HCN and biofilm matrix could reduce *C. elegans* survival (41, 42, 80).

We propose the following for host-pathogen iron competition, where intestinal *P. aeruginosa* downregulated AlgR, which led to increased expression of PvdA. Correspondingly, *C. elegans* upregulated ferritin protein that participated in iron competition with pyoverdine. Intercepting host-pathogen iron competition by inhibiting pyoverdine synthesis will tilt the favor against *P. aeruginosa* survival, thereby enabling better survival of the animal host. We identified and evaluated Galangin as a novel PvdA inhibitor, which could be used in combinatorial therapy with ciprofloxacin to effectively eradicate *P. aeruginosa* intestinal and skin infection. Galangin is used to kill cancer cells and kill *Staphylococcus aureus* at higher concentrations than the concentration used in our project (81, 82), indicating that Galangin is safe for anti-virulence purposes against *P. aeruginosa* infections. Future structure-activity relationship (SAR) studies may enhance Galangin effectiveness of a PvdA inhibitor (83). Hence, advances in ‘omics’ approaches enable concurrent study of host and pathogen interactions, where the bacterial targets can be used to develop novel antimicrobial strategies against difficult-to-treat infections.

## Method and Materials

### Growth and Maintenance of Bacterial Strains and *C. elegans*

The bacterial strains and *C. elegans* used in this project were described in **Supplementary Table 1**. Medium for bacterial cultivation was Luria-Bertani (LB) broth (Difco, BD Company, USA), while *P. aeruginosa* strains were grown in ABTGC (ABT minimal medium supplemented with 2 g L^−1^ glucose and 2 g L^−1^ casamino acids at 37 °C (84). For plasmid maintenance in *E. coli*, the medium was supplemented with 100 μg ml^−1^ ampicillin and 15 μg ml^−1^ gentamicin. For marker selection in *P. aeruginosa*, 30 μg ml^−1^ gentamicin and 100 μg ml^−1^ streptomycin were used.

Nematode growth medium (NGM) agar and liquid worm assay medium(67) were used for *C. elegans* maintenance and experiments(85). *C. elegans* used in this project were collected from the Caenorhabditis Genetics Center (CGC), the University of Minnesota. For animal cultivation, *E. coli* OP50 bacterial lawn were cultivated on NGM agar plates at 37 °C for 16 hrs. The *C. elegans* was transferred to the NGM agar plates with OP50 lawn and cultivated at room temperature for 72 hrs to allow the population expansion. For the *C. elegans ftn-2*(ok404) mutant, the NGM agar containing 5 mg ml^-1^ ammonium ferric citrate was used as cultivation medium. Since standard infection assays typically use a range of 15 – 40 nematodes per plate for experiments (86–88), at least 30 nematodes per replicate plate, unless otherwise stated were used for our study.

### SILAC Proteomics Assay

The *P. aeruginosa* lysine auxotrophic strain Δ*lysA* was grown in the liquid ABTG medium with Amino acid Drop-out Mix Minus Lysine without Yeast Nitrogen Base powder (United States Biological, MA) and 500 μM ^13^C-_L_-lysine (_L_-lysine:2HCl, U-13C6, 98%, CIL, Inc, USA) labeled overnight. The overnight culture was filtered through the 0.22 μm filter (GTTP04700, Merck Millipore, America) by the vacuum suction filtration pump to collect the bacterial cells on the filters. The filters were placed on the ABTG agar with Amino acid Drop-out Mix Minus Lysine without Yeast Nitrogen Base powder and 500 μM ^13^C-_L_-lysine to enable biofilm lawn growth for 48 hrs at 37 °C. The filter containing the biofilm lawn were next transferred to new NGM-ml agar, NGM agar minus lysine adding Amino acid Drop-out Mix Minus Lysine without Yeast Nitrogen Base powder and ^12^C-_L_-lysine additive (_L_-lysine powder, ≥ 98%,

Sigma-Aldrich, USA), where around 100 live L3 *C. elegans* animals pre-labelled with ^12^C-_L_-lysine were transferred from feeding plates to biofilm on filters. After 6 h.p.i and 24 h.p.i at 25 °C, all live nematodes were collected into 1.5 ml microcentrifuge tubes and washed by ice-cold PBS buffer via centrifugation for 10 times to remove bacteria retaining on the animal surface. Dead nematodes were not collected for proteomics. For the nematode samples, they were grinded to tissue fragments first before adding the ice-cold lysis buffer. The lysate was then sonicated and centrifuged to remove cellular debris pellet. Lysate protein concentration was determined by Pierce BCA Assay (Thermo Fisher Scientific, Franklin, Massachusetts) as per the manufacturer’s instructions. Total protein was reduced with 10 mM 1,4-dithiothreitol (DTT) at 37 °C for 1 h and subsequently alkylated in 20 mM iodoacetamide (IAA) for 30 min at room temperature in the dark. Proteins were digested with Lys-C (1:100, w/w) at 37 °C for 2 h and then trypsin (1:100, w/w) at 37 °C overnight. All the tryptic peptides samples were desalted on a Sep-pak C18 cartridges column and then lyophilized under vacuum.

The vacuum-dried samples were resuspended in 0.1% FA for liquid chromatography (LC)-MS/ MS analysis. Each sample of peptides was loaded onto a C18 trap column (75 μm ID × 2 cm, 3 μm, Thermo Scientific) and then separated on a C18 analytical column (75 μm ID×50 cm, 2 μm, Thermo Scientific). Peptides were separated and analyzed on an Easy-nLC 1200 system coupled to a Q-Exactive HF-X Hybrid Quadrupole-Orbirap Mass spectrometer system (Thermo Fisher Scientific). Mobile phase A (0.1% formic acid in 2% ACN) and mobile phase B (0.1% formic acid in 98% ACN) were used to establish a 60min gradient composed of 1 min of 6% B, 48min of 6-28% B, 1min of 28-60%B, 1 min 60-90% B, 9 min of 90% B at a constant flow rate of 250 nl/min at 55℃. MS data were acquired using the following parameters: for all experiments, the instrument was operated in the data dependent mode. Peptides were then ionized by electrospray at 2.2 kV. Full scan MS spectra (from m/z 375 to 1500) were acquired in the Orbitrap at a high resolution of 60,000 with an automatic gain control (AGC) of 3×10^6^ and a maximum fill time of 20ms. The twenty most intense ions were sequentially isolated and fragmented in the HCD collision cell with normalized collision energy of 27%. Fragmentation spectra were acquired in the Orbitrap analyzer with a resolution of 15,000. Ions selected for MS/MS were dynamically excluded for a duration of 30s.

The raw data were processed and searched using Proteome Discoverer software (PD) (version 2.2.0.388). A *C. elegans* protein database (4306 entries, updated on 11-2019), a *P. aeruginosa* protein database (55,063 sequences, 17,906,244 residues, updated on 11-2019) and the database for proteomics contaminants from MaxQuant (298 entries) were used for database searches. Semi-trypsin was set as the enzyme, and two missed cleavage sites of trypsin were allowed. Mass error was set to 10 ppm for precursor ions and 0.02 Da for fragmented ions. Carbamidomethylation on Cys was specified as the fixed modification, and oxidation (M), acetylation (Protein N-term) and isotope labeling of lysine (+6.020 Da) were set as variable modifications. Reversed database searches were used to evaluate false discovery rate (FDR) of peptide and protein identifications. False discovery rate (FDR) thresholds for protein, peptide and modification site were specified at 1%. Minimum peptide length was set at 7. All other parameters were set to default values. We used ANOVA (Background Based) tests to formalize p-value calculation and perform a multiple hypothesis testing correction on significantly different proteins, where the adjusted p-value was processed via Bonferroni-Holm method. Then the fold-change values and regulation situations of proteins of samples from 6 h.p.i to 24 h.p.i in bacterial and host groups were calculated and performed basing on the formulas respectively:

For bacteria proteins: 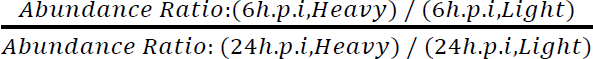

For host animal proteins: 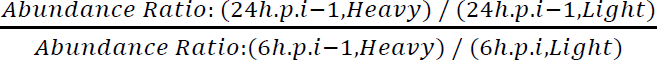

The ratio was set with the 2-fold changes and p-value <0.05. All the statistical calculation and data process were processed via MetaboAnalyst 5.0 and analysis in Language R. LC– MS/MS raw data of all replicates, results for protein and peptide identification and quantification from MaxQuant has been submitted to the ProteomeXchange Consortium as an integrated proteome resource (89).

### Quantification of Bacterial Numbers in *C. elegans* Intestine by Colony Forming Unit (CFU)

After cultivation on the NGM plates with bacteria lawn, nematodes were washed 3 times with sterile 0.9% (w/v) NaCl saline by centrifuging at 1000-2000 rpm for 30 sec. The nematodes were resuspended in 100 μl sterilized 0.9% NaCl saline in 1.5 ml-microcentrifuge tubes (JET BIOFIL, China), followed by homogenization with pellet pestles. The homogenates were serially diluted in 0.9% NaCl in 95-well plate (SPL Life Sciences company, South Korea) and subsequently cultivated on LB agar petri dishes at 37 °C for 16 hrs. After enumeration of colonies grown on agar plate, the CFU ml^-1^ were calculated by the following formula: colony number × dilution factor / volume. Experiments were performed in technical triplicates with 3 independent trials, and the results are shown as mean ± s.d.

### Structure-based virtual screening for PvdA inhibitors

Software AutoDock Vina was employed to conduct molecule docking of PvdA protein active site to the chemical library *in silico*. Discovery studio 2016 Client was used to edit the PvdA protein by eliminating water molecules and adding polar hydrogens. All ligands’ libraries were obtained from ZINC database. The OpenBabel GUI was used to convert the libraries into single ligand molecules. As reference ligand, N∼5∼-hydroxy-L-ornithine (ONH for short) was used to detect primary bonding sites on PvdA protein. Each output result included 10 models which were evaluated and ranked by binding affinity energy. The 3D images of binding models were generated by PyMOL v.2.3.2, and PyMOL was also used to select the numbers of bonds to protein active site.

### Inhibition and half maximal inhibitory concentrations (IC50) testing

To evaluate the inhibitory effect of compounds on PvdA expression, the PAO1/p*_pvdA_-gfp* was cultivated in ABTGC medium containing a range of Galangin concentrations (0–200 μM) in triplicate wells of a 96-well plate (SPL Life Sciences company, South Korea). The 96-well plate was incubated at 37 °C for 16 hrs in a microplate reader (Tecan Infinite M1000 Pro, Switzerland) for analysis of OD600 (Absorbance at wavelength 595 nm). GFP (Ex: 495 nm, Em: 515 nm) and pyoverdine (Ex: 400 nm, Em: 450 nm) every 15 mins. The half-maximal inhibitory concentration (IC50) was calculated by GraphPad Prism 8.0.2 software (GraphPad Software Inc., USA). Experiments were performed in technical triplicates with 3 independent trials, and the results were shown as the mean ± s.d.

### *C. elegans* killing Assay

*P. aeruginosa* strains were cultivated on NGM agar plates at 37 °C for 16 hrs, followed by transfer of 30 individuals/ plate of L3 *C. elegans* from OP50 feeding plate to *P. aeruginosa* lawn by titanium wire picker. Live and dead animals were enumerated daily to evaluate the killing assay of *C. elegans*.

### Galangin-ciprofloxacin combinatorial treatment against *C. elegans* intestinal infection

The infected *C. elegans* were treated with monotherapy or combinatorial therapy of Galangin and ciprofloxacin in a liquid worm assay medium (67) for 7-8 hrs daily for 156 hrs (6 days). After drug treatment, the live and dead *C. elegans* were enumerated. Experiments were performed in technical triplicates with 3 independent trials, and the results are shown as mean ± s.d.

### Epifluorescence microscopy

For epifluorescence microscopy of intestinal bacteria in the nematodes, the live nematodes were first collected and then fixed in μ-Slide 8-well plates (Ibidi, Germany) containing 2% agarose + 4% paraformaldehyde (Sigma Aldrich, USA). All microscopy images with Z-stack were captured and acquired by using Nikon Eclipse Ti2-E Live-cell Fluorescence Imaging System with 20× objective using Brightfield, GFP (Ex: 495 nm, Em: 515 nm) and pyoverdine (Ex: 400 nm, Em: 450 nm) fluorescent channels. As autofluorescence of the entire nematode body across a wide range of fluorescent spectrum poses a problem for microscopy (90), it is important to ensure that only bacterial fluorescence within the intestine were observed and measured. At least 5 images were captured for every sample. All the images were exported by the NIS (Nikon) program with 100 μm scale bars in all images.

### Tabulation of relative fluorescence density

Relative fluorescence levels in the images were quantified by ImageJ program using the equation: corrected total cell fluorescence (CTCF) = Integrated Density – (Area of selected cell X Mean fluorescence of background readings). Experiments were performed in technical triplicates with 3 independent trials, and results were shown as the mean ± s.d.

### Galangin-ciprofloxacin combinatorial treatment against Medaka wounds infection

All Medaka fish experiments were performed according to the guidance of the Department of Health in Hong Kong. They were authorized and approved by the City University of Hong Kong animal research ethics committee according to ARRIVE guidelines, with permit number Ref. No. A-0418. The Medaka fish cultivation and the wounds infection on fishes were processed and established as previously described (79). Based on power analysis and prior infection studies (45), detection of at least 1000-fold difference in CFU is minimally feasible with 8-10 animals as sample size in each group for 3 independent trials.

The infected fishes were treated with monotherapy or combinatorial therapy of Galangin and ciprofloxacin in sterile culture water for 6 hours. These treated fishes were anesthetized by immersing the animals in 0.015 μM anesthesia tricaine mesylate (MS 222) (Merck KGaA, USA) for CFU test and fluorescence images capturing.

### Statistical analysis

Experiments were performed in technical triplicates with 3 independent trials. The results were expressed as means ± standard deviation. All data sets were evaluated using the one-way ANOVA and Student’s t-test to study the associations between independent variables and calculate the *p-*values.

## Acknowledgments

This research is supported by The Hong Kong Polytechnic University, Department of Applied Biology and Chemical Technology Startup Grant (BE2B), State Key Laboratory of Chemical Biology and Drug Discovery Fund (1-BBX8), Departmental General Research Fund (UALB), One-line account (ZVVV), Environmental and Conservation Fund (ECF-48/2019), Health and Medical Research Fund (HMRF-201903032), Gifted Education Fund (2020–06) and Pneumoconiosis Compensation Fund Board (ZJN2).

We sincerely thank Prof Haihua Liang, Northwest University, for the generous gift of the ΔalgR mutant. The C. elegans strains were provided by the Caenorhabditis Genetics Center (CGC), which is funded by NIH Office of Research Infrastructure Programs (P40 OD010440).

## Author Contributions

S.L.C designed methods and experiments. Y.S.L and C.Z performed laboratory experiments, analysed the data, and interpreted the results. P.H analysed the data. S.L.C and Y.S.L, wrote the paper. All authors have contributed to, seen, and approved the manuscript.

## Data and materials availability

All data are available in the main text or the supplementary materials. The mass spectrometry proteomics raw data have been deposited to the ProteomeXchange Consortium (http://proteomecentral.proteomexchange.org) via the iProX partner repository with the dataset identifier PXD036121.

## Supplementary Materials

**Supplementary figure 1:**
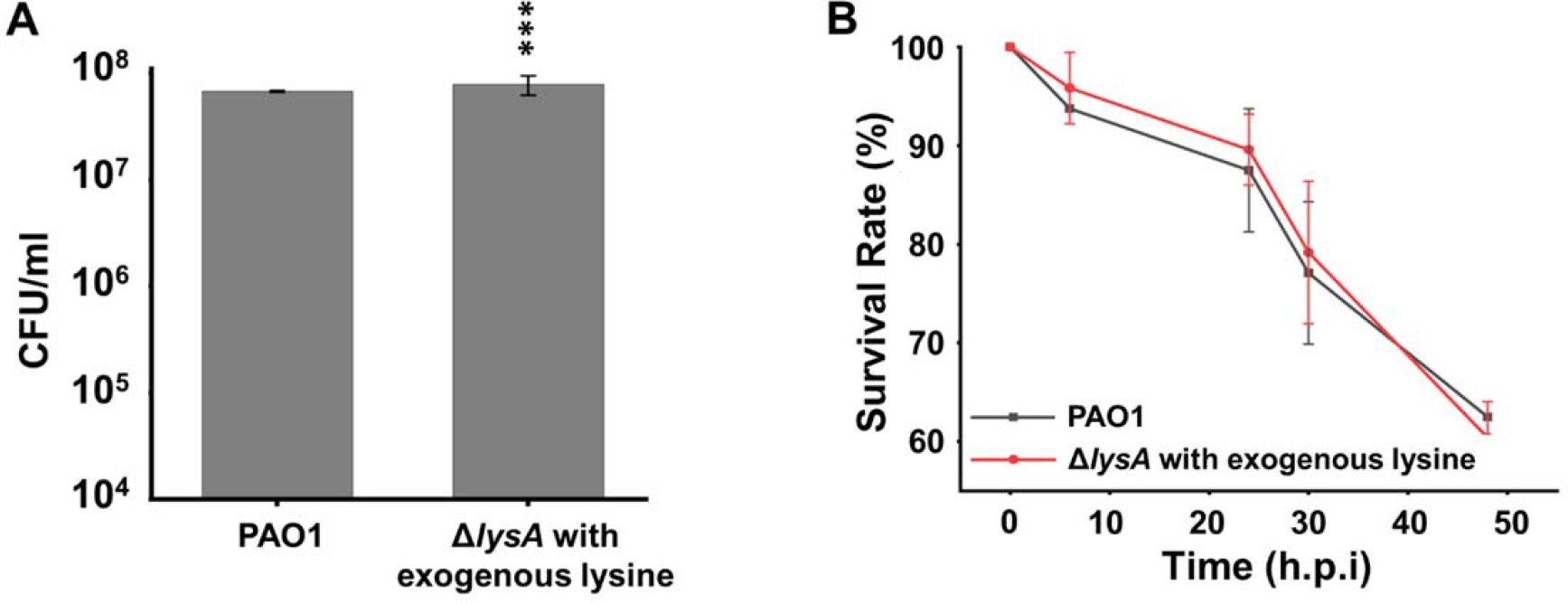
The comparison of PAO1 and mutant Δ*lysA* of *P. aeruginosa* in the *in vivo* infection to *C. elegans*. (**A**) Bacterial numbers (CFU) of PAO1 and Δ*lysA* mutant inside nematodes. (**B**) Nematode survival rate after infected by wild-type PAO1 and Δ*lysA* for days respectively. Scale bars, 100 μm. H.p.i = hours post-infection, ***P < 0.001.

**Supplementary figure 2:**
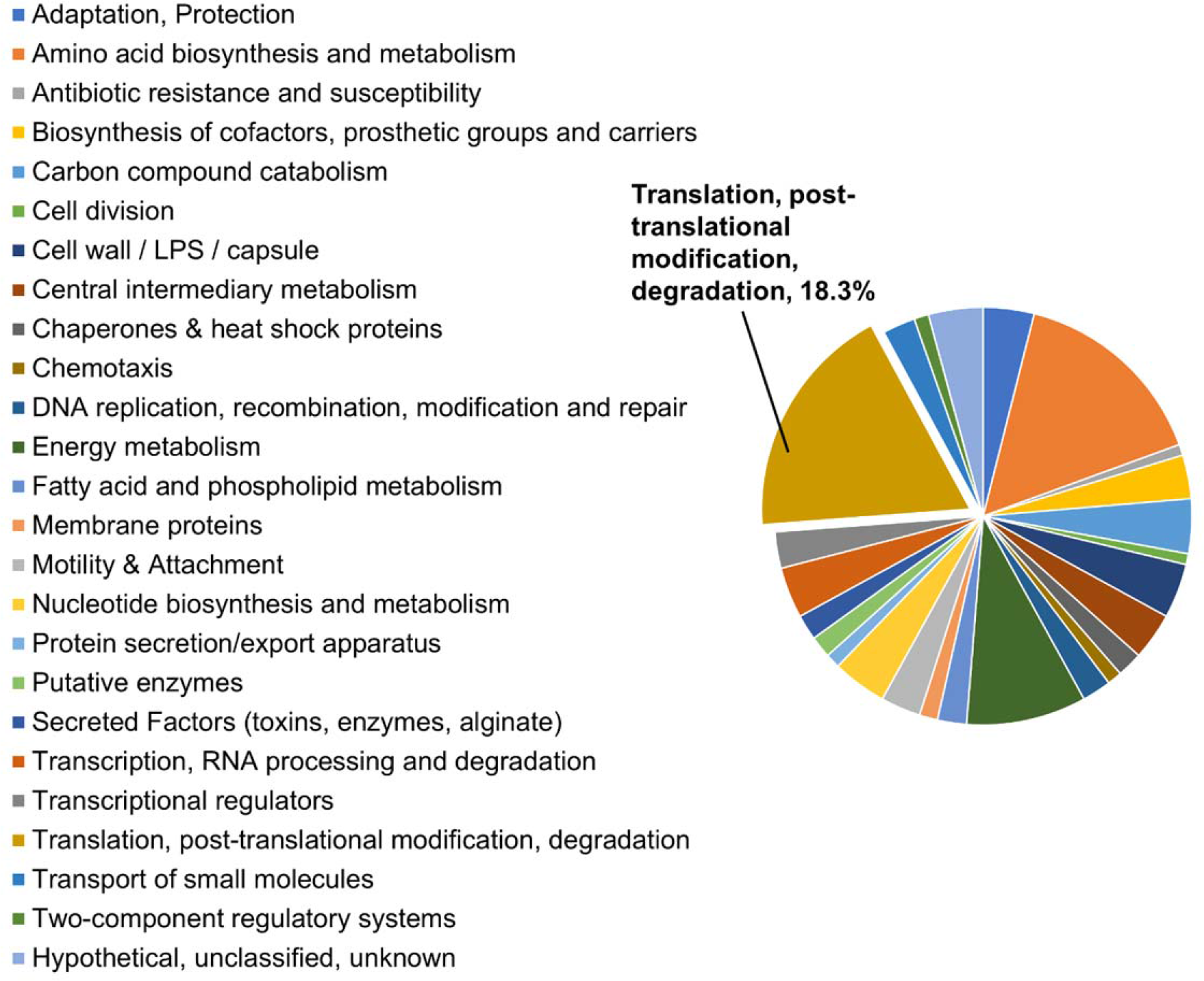
Lowly expressed functional proteins in *P. aeruginosa* during. interactions. The proteins in the functional group of Translation. Post-translational modification, degradation took the largest proportion.

**Supplementary figure 3:**
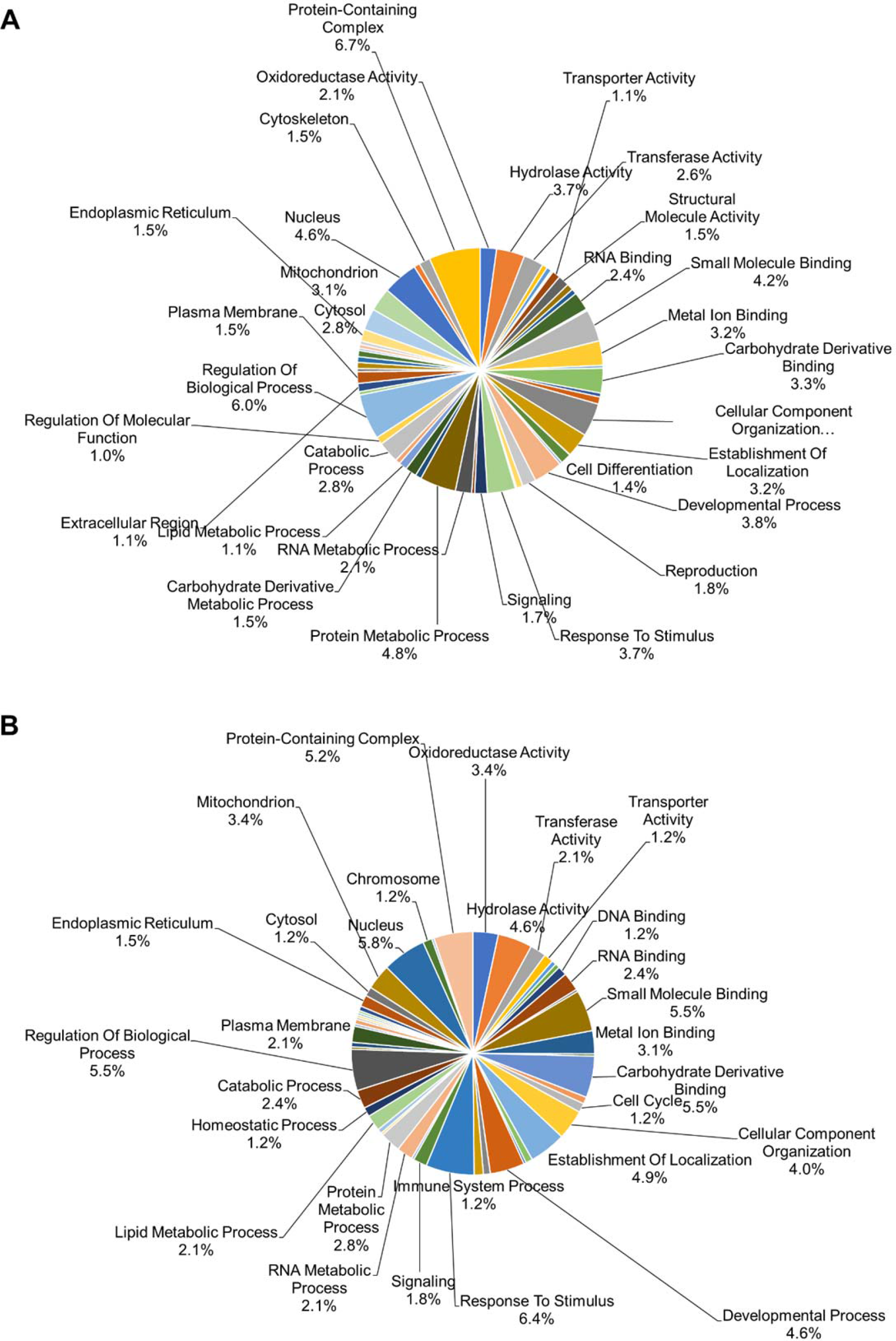
Highly and lowly expressed functional proteins in *C. elegans* during interactions. (**A**) The upregulated: high expressed proteins of the host proteins during the 18 hours interaction are in functional groups. (**B**) The downregulated proteins in *C. elegans* were classified into functional groups.

**Supplementary Figure 4:**
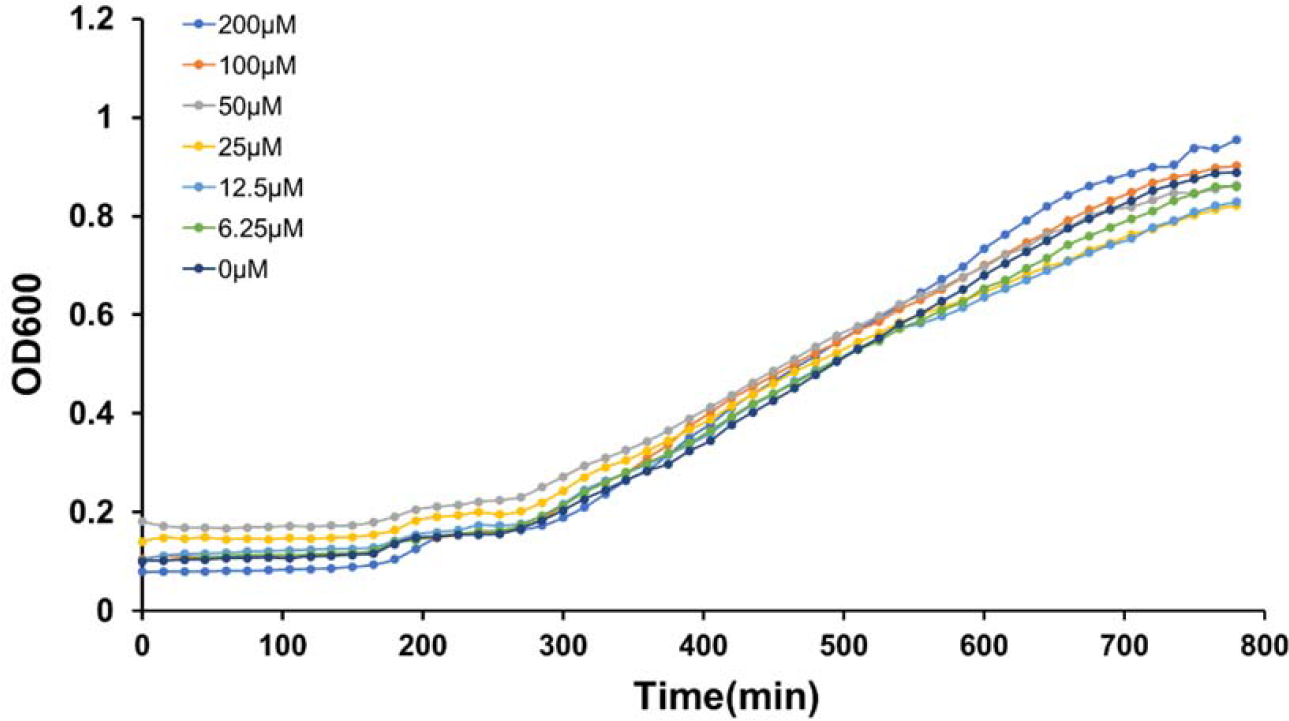
Growth curve of *P. aeruginosa* treated with Galangin (μM). Galangin could not inhibit bacterial growth over time.

**Supplementary figure 5:**
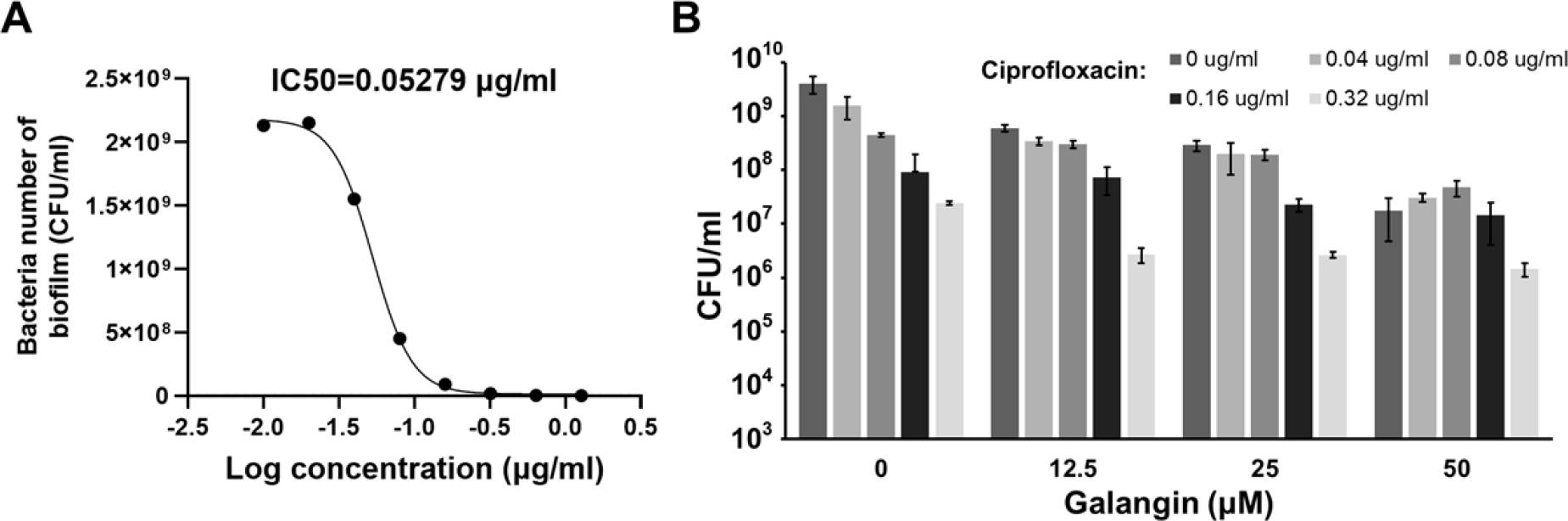
The inhibition of the biofilm formation of ciprofloxacin and Galangin in presence of the ciprofloxacin. (**A**) The IC50 of ciprofloxacin to inhibit *in vitro* biofilm formation. (**B**) The CFU/ml of *P. aeruginosa* biofilm treated by various concentrations of Galangin and ciprofloxacin *in vitro* assay. Ciprofloxacin: μg/ml, Galangin: μM.

**Supplementary table 1:**
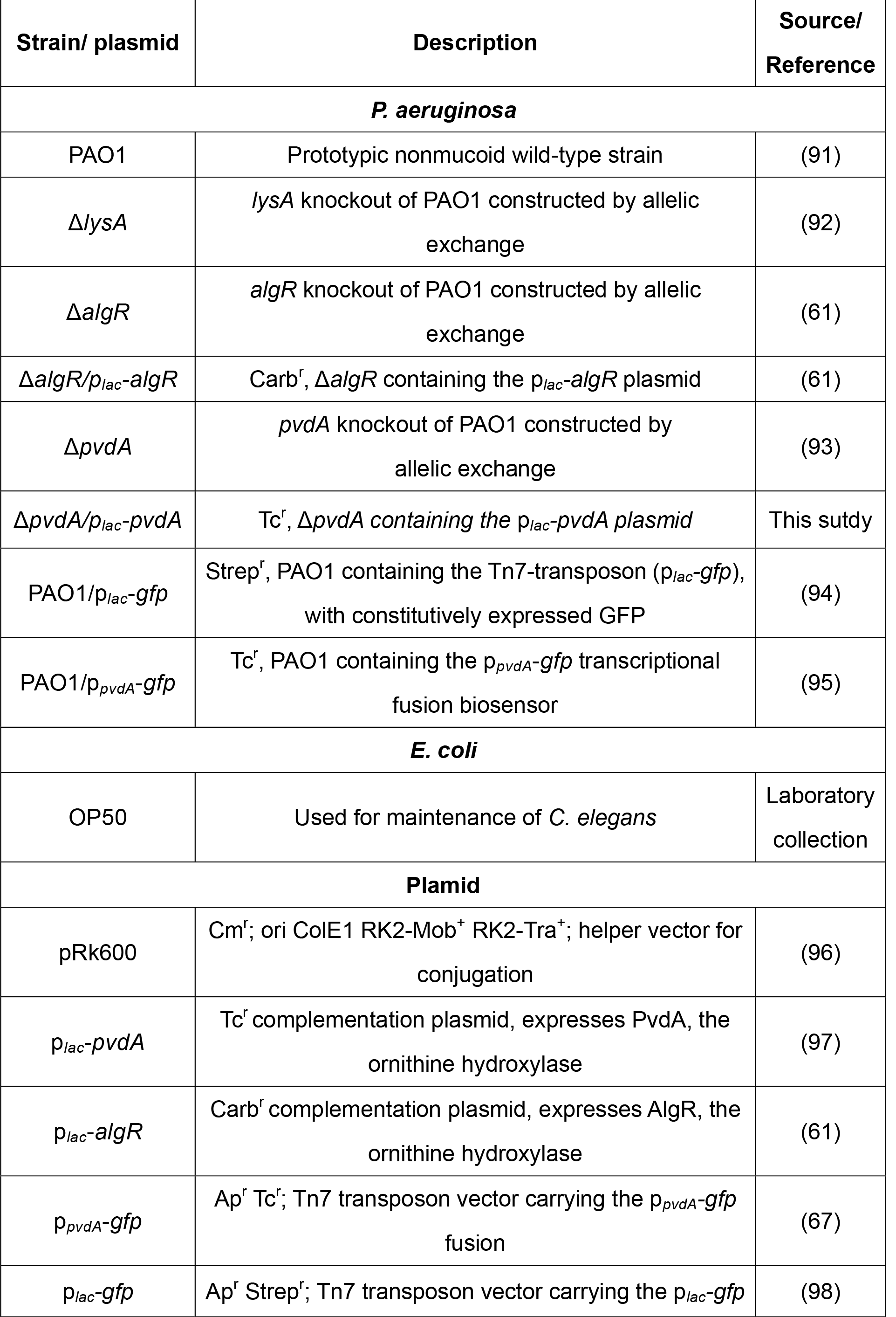
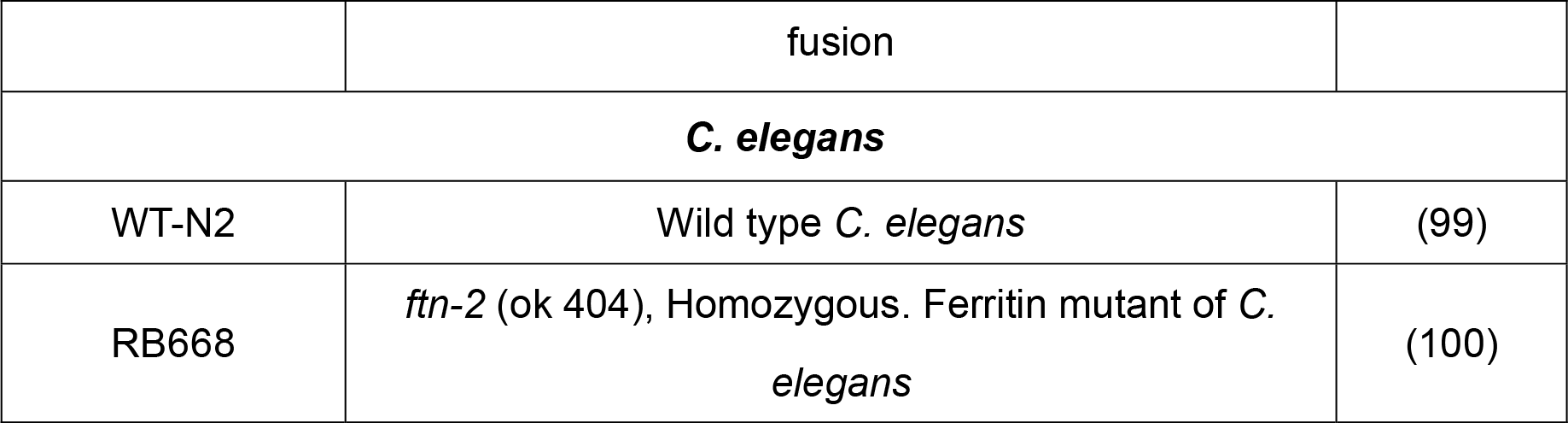
Bacterial strains, plasmids and *C. elegans* strains used in this study.

| Strain/ plasmid | Description | Source/<br>Reference |
| --- | --- | --- |
| ***P. aeruginosa*** |  |  |
| PAO1 | Prototypic nonmucoid wild-type strain | (91) |
| $\Delta lysA$ | *lysA* knockout of PAO1 constructed by allelic exchange | (92) |
| $\Delta algR$ | *algR* knockout of PAO1 constructed by allelic exchange | (61) |
| $\Delta algR/p_{lac}-algR$ | Carb<sup>r</sup>, $\Delta algR$ containing the *p<sub>lac</sub>-algR* plasmid | (61) |
| $\Delta pvdA$ | *pvdA* knockout of PAO1 constructed by allelic exchange | (93) |
| $\Delta pvdA/p_{lac}-pvdA$ | Tc<sup>r</sup>, $\Delta pvdA$ containing the *p<sub>lac</sub>-pvdA* plasmid | This study |
| PAO1/*p<sub>lac</sub>-gfp* | Strep<sup>r</sup>, PAO1 containing the Tn7-transposon (*p<sub>lac</sub>-gfp*), with constitutively expressed GFP | (94) |
| PAO1/*p<sub>pvdA</sub>-gfp* | Tc<sup>r</sup>, PAO1 containing the *p<sub>pvdA</sub>-gfp* transcriptional fusion biosensor | (95) |
| ***E. coli*** |  |  |
| OP50 | Used for maintenance of *C. elegans* | Laboratory collection |
| **Plasmid** |  |  |
| pRk600 | Cm<sup>r</sup>; ori ColE1 RK2-Mob<sup>+</sup> RK2-Tra<sup>+</sup>; helper vector for conjugation | (96) |
| *p<sub>lac</sub>-pvdA* | Tc<sup>r</sup> complementation plasmid, expresses PvdA, the ornithine hydroxylase | (97) |
| *p<sub>lac</sub>-algR* | Carb<sup>r</sup> complementation plasmid, expresses AlgR, the ornithine hydroxylase | (61) |
| *p<sub>pvdA</sub>-gfp* | Ap<sup>r</sup> Tc<sup>r</sup>; Tn7 transposon vector carrying the *p<sub>pvdA</sub>-gfp* fusion | (67) |
| *p<sub>lac</sub>-gfp* | Ap<sup>r</sup> Strep<sup>r</sup>; Tn7 transposon vector carrying the *p<sub>lac</sub>-gfp* | (98) |
|  | fusion |  |
| ***C. elegans*** |  |  |
| WT-N2 | Wild type *C. elegans* | (99) |
| RB668 | *ftn-2* (ok 404), Homozygous. Ferritin mutant of *C. elegans* | (100) |

